# Development of SARS-CoV-2 replicons for the ancestral virus and variant of concern Delta for antiviral screening

**DOI:** 10.1101/2022.10.11.511804

**Authors:** Maximilian Erdmann, Maia Kavanagh Williamson, Tuksin Jearanaiwitayakul, James Bazire, David A. Matthews, Andrew D. Davidson

**Affiliations:** School of Cellular and Molecular Medicine, University of Bristol, BS8 1TD, UK; Department of Microbiology, Mahidol University, Bangkok, Thailand

**Keywords:** SARS-CoV-2, reverse genetics, replicon, Variant of Concern, antiviral testing

## Abstract

SARS-CoV-2 is the aetiologic agent of COVID-19 and the associated ongoing pandemic. As the pandemic has progressed, Variants of Concern (VOC) have emerged with lineage defining mutations. Using a SARS-CoV-2 reverse genetic system, based on transformation associated recombination in yeast, a series of replicons were produced for the ancestral Wuhan virus and the SARS-CoV-2 VOC Delta in which different combinations of the Spike, membrane, ORF6 and ORF7a coding sequences were replaced with sequences encoding the selectable marker puromycin N-acetyl transferase and reporter proteins (*Renilla* luciferase, mNeonGreen and mScarlet). Replicon RNAs were replication competent in African green monkey kidney (Vero E6) derived cells and a range of human cell lines, with a Vero E6 cell line expressing ACE2 and TMPRSS2 showing much higher transfection efficiency and overall levels of *Renilla* luciferase activity. The replicons could be used for transient gene expression studies, but cell populations that stably maintained the replicons could not be propagated. Replication of the transiently expressed replicon RNA genomes was sensitive to remedesivir, providing a system to dissect the mechanism of action of antiviral compounds.

## Introduction

Severe acute respiratory syndrome coronavirus 2 (SARS-CoV-2) is the third and most recent highly pathogenic human coronavirus (CoV) to emerge in the last two decades (1-4). Like SARS-CoV and Middle Eastern respiratory syndrome (MERS)-CoV, it is an enveloped RNA virus that belongs to the *Betacoronavirus* genus of the *Coronaviridae* family (1). SARS-CoV-2 is the causative agent of the ongoing ‘coronavirus disease 19’ (COVID-19)-pandemic (5). The first detected case of COVID-19 appeared in Wuhan, China (December 2019) and was followed by rapid dissemination, resulting in the declaration of the current COVID-19 pandemic in March 2020 (5, 6).

Like all coronaviruses, SARS-CoV-2 has a capped, positive-sense RNA genome of ~30 kb that contains two large 5′ open reading frames (ORFs) followed by several, smaller 3′ ORFs (7). At the 5′-end of the genome there is a highly structured untranslated region (UTR) followed by ORF1a and ORF1ab that encode the large polyproteins 1a (pp1a) and pp1ab respectively (2, 8). Translation of ORF1ab is mediated by a −1 ribosomal frameshift event during translation of ORF1a and determines the ratio of ORF1a:ORF1ab encoded viral proteins (9). pp1a and pp1ab are post-translationally processed by viral encoded proteases to yield 16 non-structural proteins (nsp1-16) which are highly conserved between coronaviruses (10, 11). ORF1a and ORF1ab are followed by several smaller 3′-terminal ORFs (Figure 1A). The 3’ ORFs are each preceded by a body transcription-regulatory sequence (TRS-B). During negative strand RNA synthesis, the viral RNA polymerase halts after synthesis of the negative sense TRS-B sequence. These are complementary in sequence to the TRS located in the 5′-leader sequence (TRS-L) of the 5′ UTR, allowing for template switching during negative strand synthesis. This results in the generation of nine sub-genomic negative-sense RNA templates by discontinuous transcription which are in turn used to transcribe positive sense sgRNAs (10). The sgRNAs encode structural (Spike (S), membrane (M), envelope (E) and nucleocapsid (N)) and small accessory proteins that modulate the host immune response (7, 12-14).

**Figure 1.**
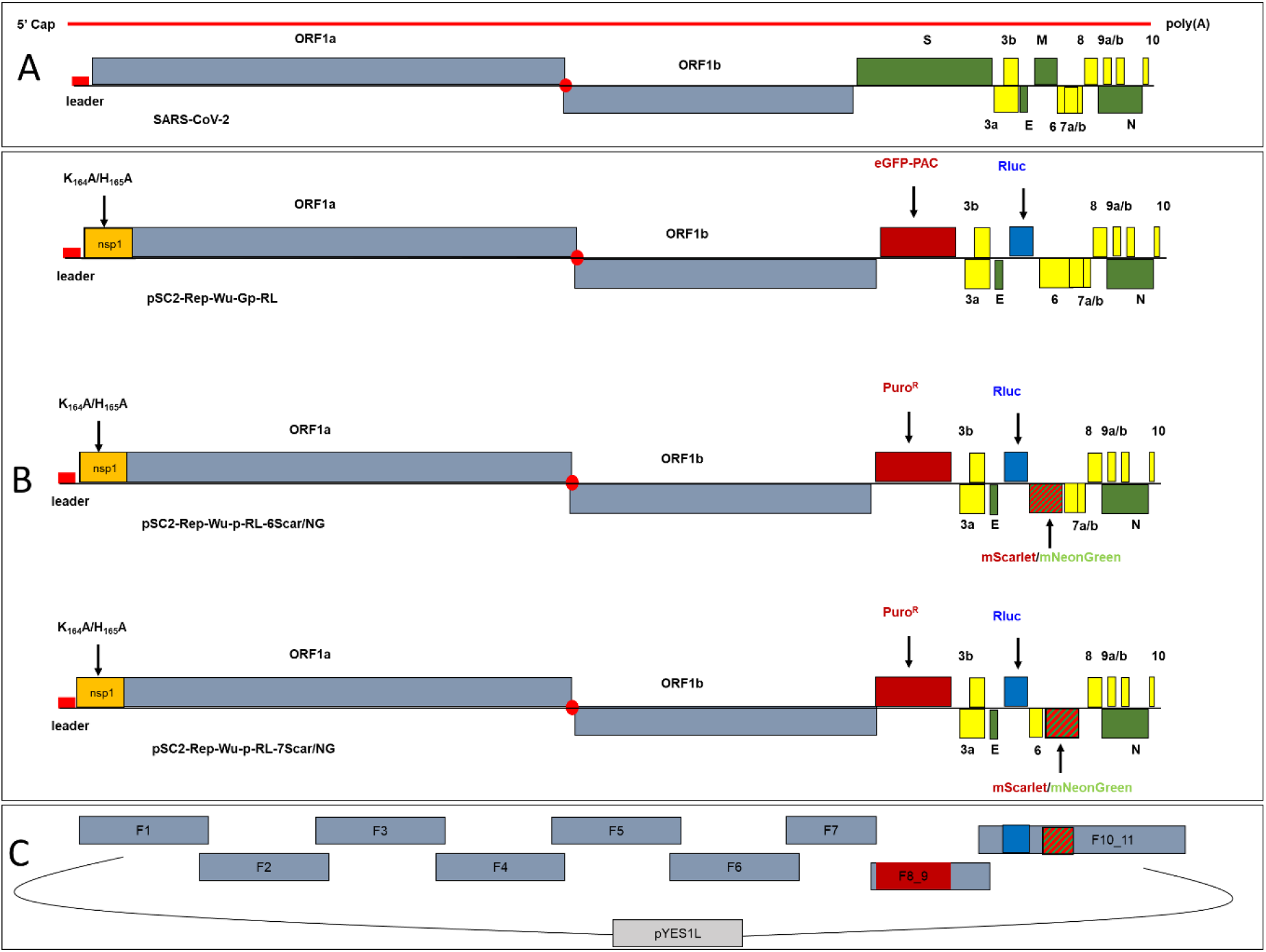
Construction of SARS-CoV-2 replicon clones. A) Schematic of the SARS-CoV-2 genome and ORFs. B) Replicon cDNA BAC/YAC constructs. For pSC2-Rep-Wu-Gp-RL, the S and M genes were replaced with sequences encoding an eGFP-pac fusion protein and RLuc respectively. Mutations encoding the amino acid changes K164A/H165A were introduced into the nsp1 sequence. pSC2-Rep-Wu-Gp-RL was modified further by replacing the eGFP-pac sequence with the *pac* gene sequence alone and ORF6 and ORF7a with either the mNeonGreen or mScarlet coding sequences, resulting in the pSC2-Rep-Wu-p-RL-6Scar/NG and pSC2-Rep-Wu-p-RL-7Scar/NG replicon clones respectively. C) Transformation associated recombination (TAR) assembly of replicon cDNA genomes. Overlapping cDNA fragments spanning the genome with 70 bp terminal end-homology were assembled into BAC/YAC shuttle vector pYES1L. Fragments F8_9 and F10_11 contain the replicon specific gene replacements.

Cellular infection is established after the SARS-CoV-2 S glycoprotein binds to the cellular angiotensin converting enzyme 2 (ACE2) receptor, initiating direct membrane fusion and/or receptor mediated endocytosis. Release of the N protein associated genome (ribonucleoprotein) into the cytoplasm initiates translation and production of the viral polyproteins. These polyproteins are co- and post-translationally cleaved into individual nsps which then assemble into viral replication complexes. Concurrently these nsps drive the distortion of the membranes of the endoplasmic reticulum-golgi intermediate complex, to form membrane enclosed vesicles within the perinuclear region (7, 9, 15).

Reverse genetics is a molecular biology technique that can be used to edit and engineer nucleic acid sequences to study the effect of a specific genotype on a phenotype (16). For viruses with positive-sense RNA genomes, the viral genome can be manipulated indirectly by cloning the full-length genome as cDNA under the control of an appropriate promoter (most commonly a T7/SP6 RNA polymerase promoter or cytomegalovirus immediate early (CMVie) promoter). The production of viral RNA transcripts with an authentic 5′-end can either occur by transcription *in vitro* (RNA launched) or can occur in host cells using host RNA polymerase II (DNA launched) such as in the case of the CMVie promoter (17, 18). In the former case the *in vitro* RNA transcripts are introduced into permissive host cells by transfection to recover virus.

For many viruses; especially highly pathogenic viruses with work restricted to biosafety level (BSL)-3 and -4, sub-genomic viral replicons have been useful surrogates to study virus replication and its effect on the host cell, as well as a valuable tool for screening antiviral compounds (17, 19-21). They may also be of particular use where viruses are not permissive to growth in standard cell culture models; as has been the case for hepatitis C virus (22). Common to all replicons is their inability to assemble infectious particles; most commonly achieved by replacing one or more structural genes with gene sequences encoding reporter proteins or selectable markers, whilst retaining all sequence motifs and gene products essential for all other aspects of the viral life cycle (most importantly genome replication) (22, 23).

The groundwork for CoV replicon systems stems from the early 2000s, with the establishment of reverse genetics systems for a number of CoVs (24-27). The first CoV replicon system was developed in 2004 for HCoV-229E, shortly after the emergence of the highly pathogenic coronavirus, SARS-CoV (16, 28). Previously sub-genomic CoV replicons, both DNA and RNA launched, have been developed using the minimal requirements for replication; the 5′/3′ UTRs and ORF1ab (16, 17, 29). The presence of the N protein acts to enhance replication and its production has been achieved both *cis*, by including the N-TRS and N coding sequence in the replicon, and *in trans*, using heterologous expression (30). Such minimal replicons allow the study of only the minimal set of viral genes for replication and when studying other aspects of the virus life cycle, e.g. particle formation or the role of accessory proteins, the replicon must be expanded to incorporate other regions of the CoV genome (17). As the introduction of replicon genomes into cells (large DNA plasmids/*in vitro* transcripts) is technically demanding and relatively inefficient, dispensable viral gene coding sequences, most commonly S, have been replaced with the coding sequence of an antibiotic resistance gene in order to select cell populations that stably express the replicon, which can then be used to analyse compounds or host-virus interactions (29, 31, 32).

To date several approaches have been undertaken for the construction of SARS-CoV-2 replicons. So far, the reported replicon systems for SARS-CoV-2 mainly relied on transfection of *in vitro* RNA transcripts (derived from replicon cDNA clones under the control of a T7 RNA polymerase promoter) into permissive cells, circumventing background expression of reporters (14, 33-41), and providing a convenient amplification step. These studies followed similar approaches used to establish replicon bearing cell lines for other CoVs and mainly relied on replacement of the S coding sequence with either sequence/s encoding reporter-selectable marker fusion proteins (14, 35, 38, 39) or a reporter protein, with an antibiotic resistance gene product encoded from a different sgRNA (e.g. E) (33, 37). In nearly all investigations, SARS-CoV-2 replicons were found to both exert high cytotoxicity, when introduced into cells either as minimal replicons containing only ORF1ab and essential control elements (14, 34, 35, 38, 42) or as larger replicons lacking one or more structural proteins but retaining the accessory genes (33, 37, 39).

The replicons constructed in this investigation retained most viral gene coding sequences except for those of S and M, to prevent assembly of infectious virions/VLPs, and either ORF6 or ORF7a. Others also opted for an approach that conserves most viral genes. Common to these studies and ours, was poor transfection efficiency of most cell lines tested with replicon RNA, as well as a failure to establish stable cell lines due to replicon cytotoxicity (33, 37, 39). However, we identified a Vero cell line overexpressing ACE2 and TMPRSS2 which is transfected with high efficiency with *in vitro* RNA transcripts derived from replicons corresponding to both the ancestral Wuhan sequence and the SARS-CoV-2 VOC Delta and is permissive for transient replicon RNA replication. As the replicons developed in this investigation express both a reporter enzyme and fluorescent marker protein, they can be used conveniently in reporter assays.

## Methods and Materials

### Cell lines

Human Caco-2 cells (ATCC® HTB-37™; a kind gift from Dr Darryl Hill, University of Bristol) were obtained from the American Type Culture Collection and used to produce Caco-2 cells constitutively expressing human ACE2 (Caco-2/ACE2 (43), a kind gift from Dr Yohei Yamauchi, University of Bristol). Human liver epithelial Huh7.5 cells (44) were a kind gift from Professor Mark Harris, University of Leeds). An African green monkey kidney cell line (Vero E6) modified to constitutively express the human serine protease TMPRSS2 (Vero E6/TMPRSS2 (VTN), (45)) was obtained from NIBSC, UK. Vero E6 cells and human A549 cells modified to constitutively express ACE2 and TMPRSS2 (VeroE6/ACE2/TMPRSS2 (VAT) and A549/ACE2/TMPRSS2 (A549-AT) cells respectively, (46))were a kind gift from Dr Suzannah Rihn, MRC-University of Glasgow Centre for Virus Research. All cells were maintained in Dulbecco’s Modified Eagle’s medium, containing 4.5 g/l D-glucose, and GlutaMAX™ (DMEM, Gibco™, ThermoFisher) supplemented with 1 mM sodium pyruvate (Sigma-Aldrich) and 10% (v/v) foetal bovine serum (FBS, Gibco™, ThermoFisher) except Caco2/ACE2 cells and Caco2 cells, which were maintained in Eagle’s Minimal Essential medium containing GlutaMAX™ (MEM, Gibco™, ThermoFisher) supplemented with 1 mM sodium pyruvate (Sigma-Aldrich), 10% (v/v) FBS and 0.1 mM non-essential amino acids (Gibco™, ThermoFisher). All cell lines were grown at 37 °C in a humidified incubator in 5% CO2.

### Bacteria and yeast

OneShot® Top10 Electrocomp™ *E. coli* (Invitrogen™, ThermoFisher) were used to propagate the GeneArt™ pYES1L vector (Invitrogen™, ThermoFisher) containing SARS-CoV-2 replicon cDNA clones and were grown overnight at 37 °C either on LB-agar plates or shaking in LB media supplemented with spectinomycin at 50 µg/ml and 100 µg/ml respectively. MaV203 competent yeast cells (MATα; leu2-3,112; trp1-901; his3Δ200; ade2-101; cyh2R; can1R; gal4Δ; gal80Δ; GAL1::lacZ; HIS3UASGAL1::HIS3@LYS2; SPAL10UASGAL1::URA3; Invitrogen™, ThermoFisher) were used for yeast-based transformation-associated recombination (TAR) and propagation of inserts contained in the pYES1L vector. Yeast were grown on 2% (w/v) agar plates containing yeast nitrogen base (YNB) without amino acids (6.8 g/l, Sigma-Aldrich) supplemented with yeast synthetic drop out (YSDO) medium supplement without tryptophan (1.92 g/l, Sigma Aldrich) and 2 % (w/v) D-(+)-glucose (YSM-Trp plates) at 30 °C.

### Generation of SARS-CoV-2 cDNA fragments for TAR

A number of approaches were used to produce SARS-CoV-2 cDNA fragments for the assembly of replicon cDNA clones by TAR as follows.

a. **Synthetic DNA**. Initially nine cDNA fragments with 70 bp end-terminal overlaps were used to assemble a SARS-CoV-2 replicon clone based on the Wuhan-Hu-1 genome sequence (GenBank accession: NC_045512). The cDNA fragments were produced by GeneArt™ synthesis (Invitrogen™, ThermoFisher) as cDNA inserts in sequence verified, stable plasmid clones. The 5′ terminal cDNA fragment was modified to contain 70 nucleotides corresponding to nucleotides (nts) 9311-9380 of the pYes1L vector, a T7 RNA polymerase promoter and an extra “G” nucleotide immediately upstream of the SARS-CoV-2 5′-terminal genome sequence, whilst the 3′-terminal cDNA fragment was modified such that the 3′ end of the SARS-CoV-2 genome was followed by a stretch of 33 “A”s followed by the unique restriction enzyme site *Asc*I and nts 1-70 of the pYES1L vector. The first seven cDNA fragments (from the 5′end) spanned nts 1 – 20,090 of the SARS-CoV-2 genome. The two remaining cDNA fragments spanned nts 20,021 – 29,903 of the SARS-CoV-2 sequence with the following exceptions; nts 21,653 – 25,384, encoding the S protein, were replaced with a 1,359 nt sequence encoding an eGFP-puromycin N-acetyl transferase (pac) fusion protein and nts 26,523 – 27,191 encoding the M protein, were replaced with a 936 nt sequence encoding *Renilla* luciferase (RLuc). The cDNA fragment contained in each clone was PCR amplified using gene specific primer pairs and the Platinum SuperFi II mastermix (Invitrogen™, ThermoFisher) following the manufacturer’s instructions. The location of the fragments and primers and sequences of introduced genes are shown in Supplementary Table 1.
b. **Overlap PCR mutagenesis**. Modification of the synthetic replicon cDNA clones to introduce site-specific mutations, gene substitutions and a hepatitis delta virus ribozyme sequence followed by a T7 RNA polymerase terminator sequence (see Supplementary Table 1, a kind gift from Professor Arvind Patel, MRC-University of Glasgow Centre for Virus Research) immediately downstream of the 3′ end poly-A tail was done by overlap-PCR (OL-PCR) mutagenesis. Template DNA fragments for OL-PCR were first produced as overlapping sub-fragments (20 – 30 nt overlap) using an outer primer and an internal mutagenesis primer (primers shown in Supplementary Table 2). The first round PCR sub-fragments were purified by extraction from an agarose gel using a GeneJET Gel Extraction Kit (Thermo Scientific™, ThermoFisher). A total mass of 10-15 ng of sub-fragments at a 1:1 molar ratio were then used as templates for OL-PCR using the forward and reverse outer primers. Assembly of more than two PCR fragments was performed stepwise.
c. **Viral RNA extraction and reverse-transcriptase (RT)-PCR**. For production of a SARS-CoV-2 VOC Delta replicon, fragments 2, 4, 5 and 6 corresponding to the Wuhan-Hu-1 virus genome (Figure 1) were swapped for the corresponding cDNA fragments generated by RT-PCR from VOC Delta extracted RNA. Viral RNA was extracted from 140 µl of virus stock (SARS-CoV-2 VOC Delta, GISAID ID: EPI_ISL_15250227) using a QIAamp Viral RNA Mini Kit (Qiagen) following the manufacturer’s instructions. RT-PCR was performed using 1 µl of eluted RNA and a SuperScript™ IV One-Step RT-PCR System (Invitrogen™, ThermoFisher) as described by the manufacturer.
d. **HiFi DNA assembly**. To produce SARS-CoV-2 VOC Delta replicon clones, the viral sequence from 20,021 – 29,903 was replaced with either of two cDNA fragments in which the S and M gene coding sequences were replaced with those of the *pac* and *Rluc* genes and either the ORF6 or ORF7a coding sequences were replaced with those of the mScarlet and mNeonGreen genes respectively. Assembly of the two cDNA fragments was done using five overlapping cDNA fragments containing the VOC lineage defining mutations and replicon specific gene replacements (see Supplementary Table 3) using a NEBuilder® HiFi DNA Assembly Master Mix (NEB) according to the manufacturer’s recommendations. They were assembled in the vector pYES1L, the assembly reactions purified and electroporated into One Shot™ TOP10 Electrocomp™ E. coli. Two colonies were picked, screened for correct assembly and used as PCR templates. The resulting fragments were used for TAR assembly.

All cDNA fragments generated by PCR/RT-PCR were purified using a GeneJET PCR Purification kit (Thermo Scientific) following the manufacturer’s instructions prior to use in TAR. The concentration of purified DNA fragments was determined by spectrophotometry using a NanoDrop One (Thermo Scientific™, ThermoFisher). The regions encompassed by the cDNA inserts and the sequences of the primer pairs and 5’ and 3’ cDNA fragment modifications are shown in Supplementary Tables 1-3.

### Transformation-associated recombination (TAR) assembly in yeast

SARS-CoV-2 replicon cDNA fragments were assembled into full-length replicon cDNA clones by TAR assembly using the GeneArt™ High-Order Genetic Assembly System (Invitrogen™, ThermoFisher) according to the manufacturer’s instructions. Briefly, 200 ng of each cDNA fragment was mixed with 100 ng of the pYES1L BAC/YAC shuttle vector with Sapphire™ Technology (Invitrogen™, ThermoFisher) and the volume reduced to 5 - 10 µl using a SpeedVac™ (Savant, ThermoFisher). 100 µl of thawed MaV203 competent yeast cells were added to the tube, followed by 600 µl of PEG/LiAc. The solutions were mixed gently and incubated at 30 °C for 30 minutes, followed by the addition of 35.5 µl of dimethyl sulfoxide and incubation at 42 °C for 20 minutes. The yeast were then plated on YSM-Trp plates and grown at 30 °C until colonies were visible.

### Yeast colony screens

Correctly assembled replicon clones were identified by PCR screening of either yeast colonies directly or yeast cell lysates. For direct colony PCR, a yeast colony was picked from a YSM-Trp plate, resuspended in 100 µl of ultra-pure water and 1 µl used as a template for PCR. Yeast cell lysates were prepared following an adjusted GC preparation method (47) as follows. A yeast colony was picked from a YSM-Trp plate and resuspended in 100 µl of 5% w/v Chelex-100 (Sigma-Aldrich). Half the sample volume of acid washed glass beads were added and the sample vortexed at maximum speed for 4 minutes at room temperature. The lysate was then heated at 99 °C for 2 minutes, followed by centrifugation at 13,000 x *g* for 1 minute. 50 µl of supernatant was transferred into a new tube and 1 µl used as template for PCR. Correct assembly of the YAC clones was verified by amplifying the overlapping junctions between the fragments by multiplex PCR, using two sets of multiplex primers and the Platinum SuperFii II PCR Master Mix. The primers used in each multiplex reaction are shown in Supplementary Table 4.

### Transformation of E. coli with BAC/YAC shuttle plasmids

Correctly assembled SARS-CoV-2 replicon BAC/YAC clones were prepared for bacterial transformation as described in the GeneArt™ High-Order Genetic Assembly System protocol. An *E. coli* Pulser™ Transformation Apparatus (BioRad) was used to electroporate One Shot™ TOP10 Electrocomp™ *E. coli* in a 0.2 cm cuvette. Electroporation was performed using conditions of 2.5 kV, 200 Ω and 25 µF. Immediately after electroporation, 500 µl of S.O.C. medium (Invitrogen™, ThermoFisher) was added, and the cells incubated at 37 °C for 1 hour with shaking. Positive clones were selected by overnight growth on LA-spectinomycin (100 µg/ml) plates at 37 °C.

### BAC/YAC purification

Large-scale purification of BAC/YAC plasmid DNA was done using a NucleoBond^®^ Xtra BAC (Macherey-Nagel) kit. Single *E. coli* colonies were picked, resuspended in 2 ml of LB and used to inoculate 500 ml of LB media containing spectinomycin (50 µg/ml). After overnight growth at 37 °C with shaking, bacteria were harvested by centrifugation at 6000 x *g* for 30 minutes. Cell pellets were either frozen at −80 °C or immediately used for plasmid isolation according to the manufacturer’s protocol. The final DNA pellets were dissolved in ultra-pure water and the DNA yield and purity were determined by spectrophotometry.

### Preparation of in vitro RNA transcripts

T7 polymerase promoter driven *in-vitro* RNA transcription (IVT) from replicon BAC/YAC clones and a construct containing a codon optimised SARS-CoV-2 N gene sequence, under control of a T7 promoter (Supplementary Table 3), were performed using a RiboMAX™ Large Scale RNA Production System-T7 (Promega). When required, the BAC/YAC/plasmid constructs were linearised prior to *in vitro* transcription using a unique *Asc*I cleavage site immediately downstream of the polyA sequences. The linearised DNA constructs were then purified by phenol/chloroform extraction followed by sodium acetate/ethanol precipitation. Constructs containing a 3’-terminal hepatitis delta virus ribozyme and T7 transcription terminator were transcribed directly from purified BAC/YAC DNA. For a 50 µl reaction 1.5 µg of BAC/YAC/plasmid DNA template was incubated with 7.5 mM of ATP, CTP, UTP and 1.5 mM GTP. m7G(5’)G RNA Cap Analog (NEB) or Anti-Reverse Cap Analogue (NEB) was added to a final concentration of 3 mM followed by the addition of 5 µl of T7 Enzyme Mix and 50 U RNasin RNase Inhibitor (Promega). To produce transcripts for mock transfections the GTP concentration was increased to 7.5 mM and cap analogues were not added. The reaction mixes were then incubated at 30 °C for 3 hours. After incubation, the reaction mix was treated with RQ1 RNase-Free DNase to degrade template DNA. The reaction mix was then either used directly for electroporation or purified by LiCl precipitation, followed by three 75% ethanol washes and resuspended in ultra-pure water. Both were stored at −80 °C.

### Mammalian cell electroporation and replicon transfection

Electroporation was carried out using a Neon® Transfection System (Invitrogen™, ThermoFisher) following the manufacturer’s protocol for adherent cell lines (electroporation conditions for each cell line are shown in Supplementary Table 5). Briefly, the cells to be transfected were grown to 60-80 % confluence, detached using trypsin, harvested by centrifugation and then washed in PBS before being resuspended in resuspension buffer at a density of 1 × 10^7^ cells/ml. 110 µL of cells were then mixed with 5-15 µg of replicon *in vitro* RNA transcripts and 2 µg of N gene *in vitro* RNA transcripts in a total volume of 11 µl. Alternatively, up to 9.9 µl of replicon IVT reaction mixture was used instead of LiCl purified RNA. Immediately after electroporation, cells were resuspended in complete warm growth media. Cells were then plated at an appropriate density after resting at 37 °C for 10 minutes.

### Puromycin selection of replicon transfected cells

Selection of cells transfected with *in vitro RNA* transcripts produced from replicon constructs was initiated by addition of puromycin (InvivoGen, UK) to the culture media at 24 - 48 hours post-transfection. The optimum dose for each cell line was determined using antibiotic kill curves and selected to achieve complete death of control cells in 5 - 7 days. The drug concentrations used were 4 µg/ml for Caco2 cells and derivatives and VeroE6 cells and derivatives; 3.5 µg/ml for Huh7.5 cells and 2.5 µg/ml for A549-AT cells. Culture media with fresh drug was renewed every 3-4 days and flasks monitored for colony formation.

### Immunofluorescence assays

Cells transfected with replicon *in vitro* RNA transcripts were seeded in appropriate media in µClear 96-well Microplates (Greiner Bio-one) and grown at 37 °C in 5% CO2. After an appropriate incubation period the cells were fixed in 4 % paraformaldehyde (v/v) in PBS for 10 minutes. Fixed cells were permeabilized with 0.1% Triton-X100 in PBS and blocked with 1% (w/v) bovine serum albumin before staining with a monoclonal antibody against either the SARS-CoV-2 N protein (1:1000 dilution; 200-401-A50, Rockland), double stranded-RNA (1:250 dilution, J2 10010200, Scicons) or GFP (1:500 dilution, AB0020-200, SICGEN) followed by an appropriate Alexa Fluor-conjugated secondary antibody (1:3000 dilution, Invitrogen™, ThermoFisher) and DAPI/Hoechst 33342 (Sigma Aldrich). To determine the number of replicon expressing cells, images were acquired on an ImageXpress Pico Automated Cell Imaging System (Molecular Devices) using a 10X objective. Stitched images of 9 fields covering the central 50% of the well were analysed using Cell Reporter Xpress software (Molecular Devices).

### Luciferase assays

Assays to measure *Renilla* luciferase (RLuc) activity were done using a *Renilla* Luciferase Assay System (Promega) following the manufacturer’s recommended protocol. In short, cells were prepared and seeded in the appropriate culture vessels. To harvest the cells the media was removed and cells washed once with PBS. Lysis was achieved by incubating the cells in the recommended volume of freshly prepared 1X *Renilla* Luciferase lysis buffer for the culture vessel. Lysates were then stored at −20 °C until assayed for RLuc activity as half-scale reactions. Luciferase assays were performed in white LUMITRAC plates (Greiner Bio-one) using a GloMAX® Explorer microplate reader (Promega).

### Replicon antiviral assay

Transfected cells were seeded into 96-well plates at a density of 10,000 cells per well. An eight point; half-log fold dilution series of remdesivir (Cambridge Bioscience); starting at 20 µM (final concentration, dissolved in dimethyl sulfoxide (DMSO)) was transferred to the plates. The remdesivir concentrations/dilutions were prepared as 2X concentrates, diluting the drug to the final 1X concentration directly in each well. The final concentration of the DMSO vehicle control was 0.1% (v/v), equivalent to the DMSO concentration of the 20 µM remdesivir dose. The replicon replication initiated from *in vitro* RNA transcripts was then assayed by quantification of the RLuc signal produced in the treated cells. These values were normalised as a percentage of the vehicle control. The IC50 for remdesivir were determined using GraphPad prism (v9.4.1) by fitting a 4-parameter logistic regression curve to the data.

## Results and Discussion

A number of sub-genomic SARS-CoV-2 replicon cDNA clones were designed and tested during the investigation, with the overall aim of producing an RNA launched replicon that could be stably maintained in cells.

### Construction of a first-generation Wuhan-Hu-1 SARS-CoV-2 replicon

Initially nine cDNA fragments with 70 bp end-terminal overlaps spanning the entire SARS-CoV-2 isolate Wuhan-Hu-1 genome, except for the S and M gene coding sequences, which were replaced with sequences encoding an eGFP-pac fusion protein and RLuc respectively, were assembled into the pYES1L BAC/YAC shuttle vector using TAR assembly in yeast (Figure 1). Previously TAR has been used in the assembly of recombinant CoV genomes and replicons including in the production of the first recombinant version of SARS-CoV-2 (18, 39) The 5′-terminal cDNA fragment overlapped the pYES1L vector and contained a T7 RNA polymerase promoter and an extra “G” nucleotide immediately upstream of the SARS-CoV-2 5′ genome sequence to allow efficient transcription by T7 polymerase. The 3′-terminal cDNA fragment, which also overlapped the pYES1L vector contained a stretch of 33 “A”s at the end of the SARS-CoV-2 genome to serve as the poly-A tail, followed by the unique restriction enzyme site *Asc*I.

Mutations were introduced into the 5′-terminal cDNA fragment, encoding the nsp1 amino acid substitutions K164A/H165A, to potentially reduce cytotoxic effects induced by replicon replication. Nsp1 is the first SARS-CoV-2 protein translated and has been reported to inhibit host translation (48-50). It has low sequence conservation amongst the betacoronaviruses (and is absent in gamma-/deltacoronaviruses), however its ability to interfere with host innate immunity at the transcriptional and/or translational level is highly conserved (e.g. for murine hepatitis virus (MHV), SARS-CoV and MERS-CoV) (49-52). The nsp1 residues K164 and H165 are essential for inhibition of host cell translation (48, 53-55) and the double substitution K164A/H165A was previously reported to lead to complete loss of SARS-CoV and SARS-CoV-2 nsp1 function with respect to translation shut-off (48, 53, 54). The double substitution has been introduced into SARS-CoV-2 replicons in other investigations reducing replicon cytotoxicity (39) and leading to the recovery of cells stably harbouring a replicon (37). The resultant SARS-CoV-2 replicon was termed pSC2-Rep-Wu-Gp-RL.

*In vitro* RNA transcripts were prepared from the linearised pSC2-Rep-Wu-Gp-RL plasmid and transfected into four different mammalian cell lines to test the replicative ability of the replicon; African green monkey kidney cells (Vero E6) modified to express TMPRSS2 (VTN), Vero E6 cells modified to express TMPRSS2 and ACE2 (VAT), human lung epithelial cells (A549) modified to express TMPRSS2 and ACE2 (A549-AT) and human liver epithelial cells (Huh 7.5). *In vitro* RNA transcripts from the replicon construct were capable of initiating replication in all cell types, as determined by the detection of RLuc activity (Figure 2). Rluc is encoded by a sgRNA produced during virus replication and is therefore a marker for the replication of replicon RNA. Luminescence was detected consistently at low levels as early as 12 hours post-transfection for all cells. The kinetics of RLuc expression were similar in the four cell types with maximal RLuc activity observed between 24-36 hours post-transfection over a 96-hour observation period. However, the levels of RLuc activity were markedly different in the VAT cells (peak of 2.5 × 10^7^ relative luminescence units (RLU)) compared to the other three cell types (peaks of 1.5 – 3 × 10^5^ RLU). To investigate this further, the transfection efficiency of the four cell lines with the replicon transcripts was monitored by eGFP fluorescence. Although RLuc activity was detected in all four cell types, eGFP fluorescence was not. To determine if the eGFP-pac fusion protein was expressed by the replicon RNA transcripts, eGFP was detected by antibody staining and replicon replication examined by antibody staining for dsRNA and the N protein, both markers of coronavirus replication (7, 15). Transfection efficiency was found to be cell type dependent, with the highest transfection efficiencies identified for VAT cells, routinely ranging between 10-20 % (Figure 3).

**Figure 2.**
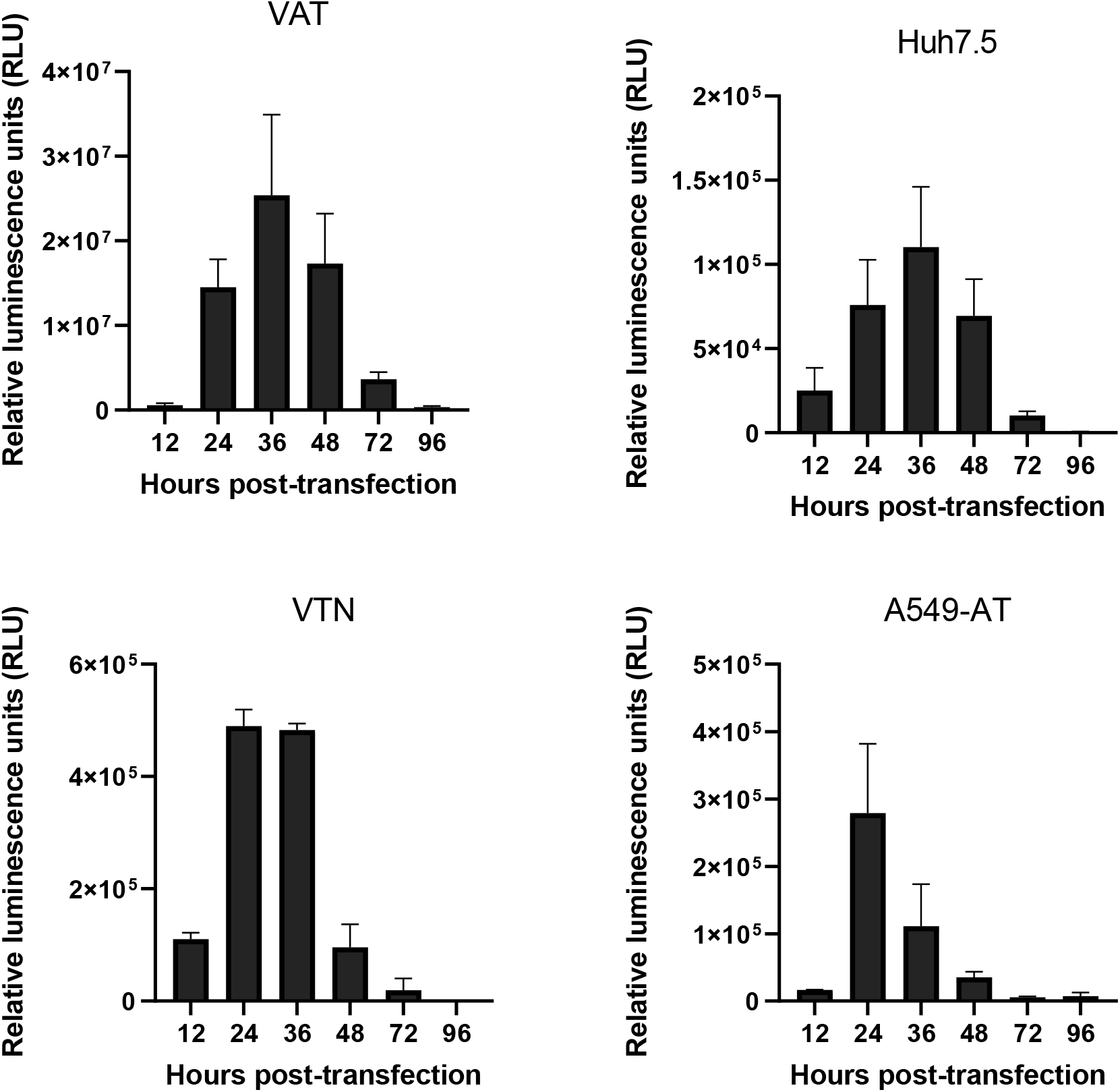
Expression of RLuc following transfection of pSC2-Rep-Wu-Gp-RL *in vitro* RNA transcripts into different cell types. VAT, Huh 7.5, VTN and A549-AT cells were transfected with *in vitro* RNA transcripts produced from the linearised pSC2-Rep-Wu-Gp-RL replicon and N gene constructs. At the indicated times post-transfection the cells were lysed and the lysates assayed for RLuc activity. The luminescence of each sample was adjusted for background by subtracting a cell only control. Graphs show the mean and standard deviation of n=4 (VAT), n=3 (Huh7.5) and n=2 (VTN, A549/A2T2) biological repeats. Each repeat was performed as a technical triplicate.

**Figure 3.**
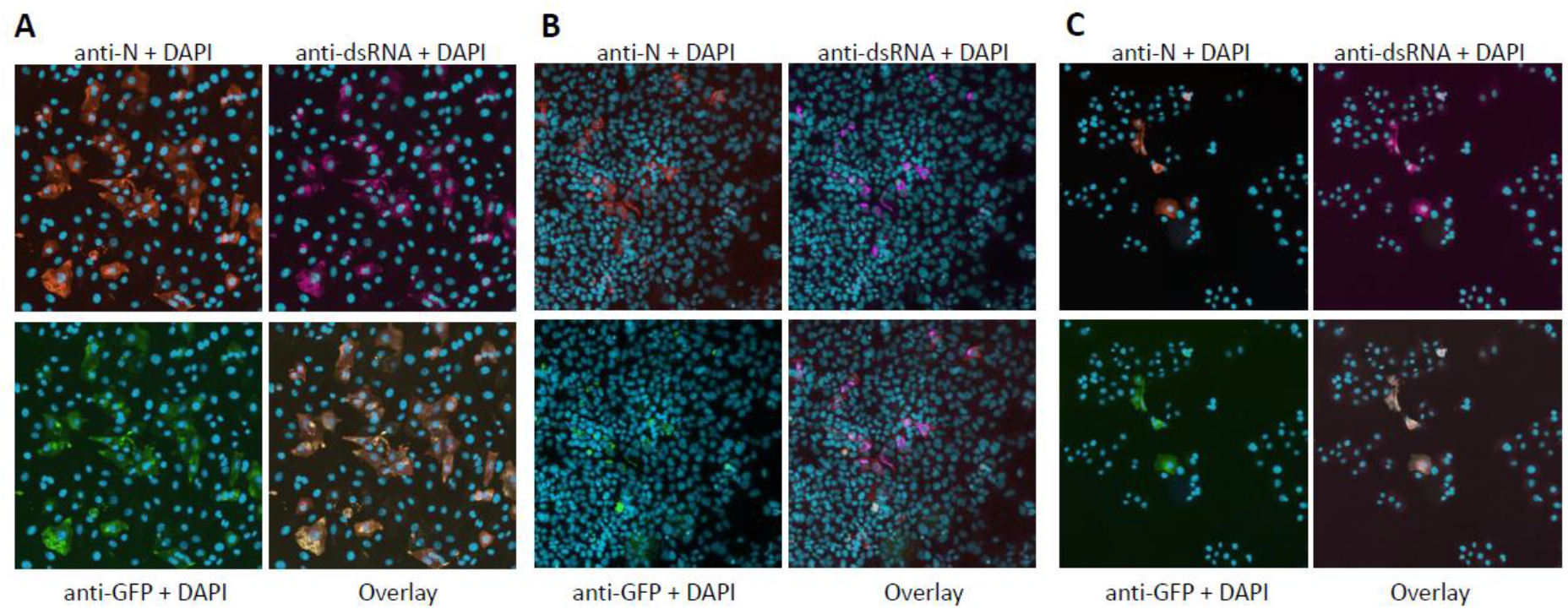
Detection of eGFP and markers of replicon replication by immunofluorescence assay. A) VAT, b) Huh 7.5 and C) VTN cells were transfected with *in vitro* RNA transcripts produced from the linearised pSC2-Rep-Wu-Gp-RL replicon and N gene constructs. Cells in the top panels were stained with antibodies recognising the SARS-CoV-2 N protein and dsRNA with nuclear DNA identified by DAPI staining. The bottom panels show staining with an antibody recognising eGFP, nuclear DNA stained with DAPI and a four-colour overlay. The number of transfected cells was determined by scoring the cells using the Cell Reporter Xpress software based on eGFP and nuclear DAPI staining of images acquired with an ImageExpressPico microscope.

Although the pSC2-Rep-Wu-Gp-RL replicon construct was useful for monitoring replicon replication, it did not express detectable levels of eGFP fluorescence and was therefore not suitable for the selection of cell lines stably expressing the replicon. Nevertheless, transfected cells were placed under puromycin selection, but it was not possible to isolate cells that proliferated under selection after 10 days. The eGFP-pac gene encodes a functional fusion protein as it was derived from a dengue virus replicon cDNA clone that was used to produce puromycin resistant and green, fluorescent cells that stably express a dengue virus replicon (56). Sequencing of the SARS-CoV-2 eGFP-pac replicon construct verified the intact sequence was present. It is likely that the eGFP-pac fusion protein was either produced at very low levels or folded improperly in cells expressing the SARS-CoV-2 replicon. Therefore, the pSC2-Rep-Wu-Gp-RL was modified further with the aim of producing a functional dual reporter replicon construct.

### Construction of second-generation Wuhan-Hu-1 SARS-CoV-2 dual-reporter replicons

To produce a functional dual reporter replicon, the Spike and M gene coding sequences were replaced with a codon-optimised *pac* gene coding sequence and RLuc coding sequence respectively. Furthermore, to produce replicons that could express a fluorescent marker protein, either the ORF6 or ORF7a coding sequences were replaced with codon optimised mNeonGreen (57) and mScarlet (58) coding sequences (Figure 1B, Supplementary Table 3). As the ORF6 TRS is located within the coding sequence of the M gene (2), which was replaced with the RLuc coding sequence, the TRS was faithfully re-introduced upstream of the ORF6 gene during the cloning process. In addition, the 3′-terminus of the replicon was modified to introduce a hepatitis delta virus ribozyme sequence and T7 RNA polymerase terminator sequence immediately downstream of the encoded poly-A tail (Supplementary Table 1), removing the need to linearise the replicon plasmids prior to *in vitro* transcription. The sequences encoding the nsp1 amino acid changes K164A/H165A were retained in the replicon clones, which were termed pSC2-Rep-Wu-p-RL-6NG, pSC2-Rep-Wu-p-RL-6Scar, pSC2-Rep-Wu-p-RL-7NG and pSC2-Rep-Wu-p-RL-7Scar (Figure 1B).

To test the functionality of the four replicon constructs, they were used to prepare *in vitro* RNA transcripts and transfected into VAT cells. At 24- and 36-hours post-transfection, the cells were fixed and examined for the presence of dsRNA and fluorescence from the mNeonGreen and mScarlet reporter proteins. *In vitro* RNA transcripts derived from all four replicon constructs were found to be replication competent, as determined by the identification of dsRNA and the expression of mNeonGreen or mScarlet from sgRNA transcripts (Figure 4).

**Figure 4.**
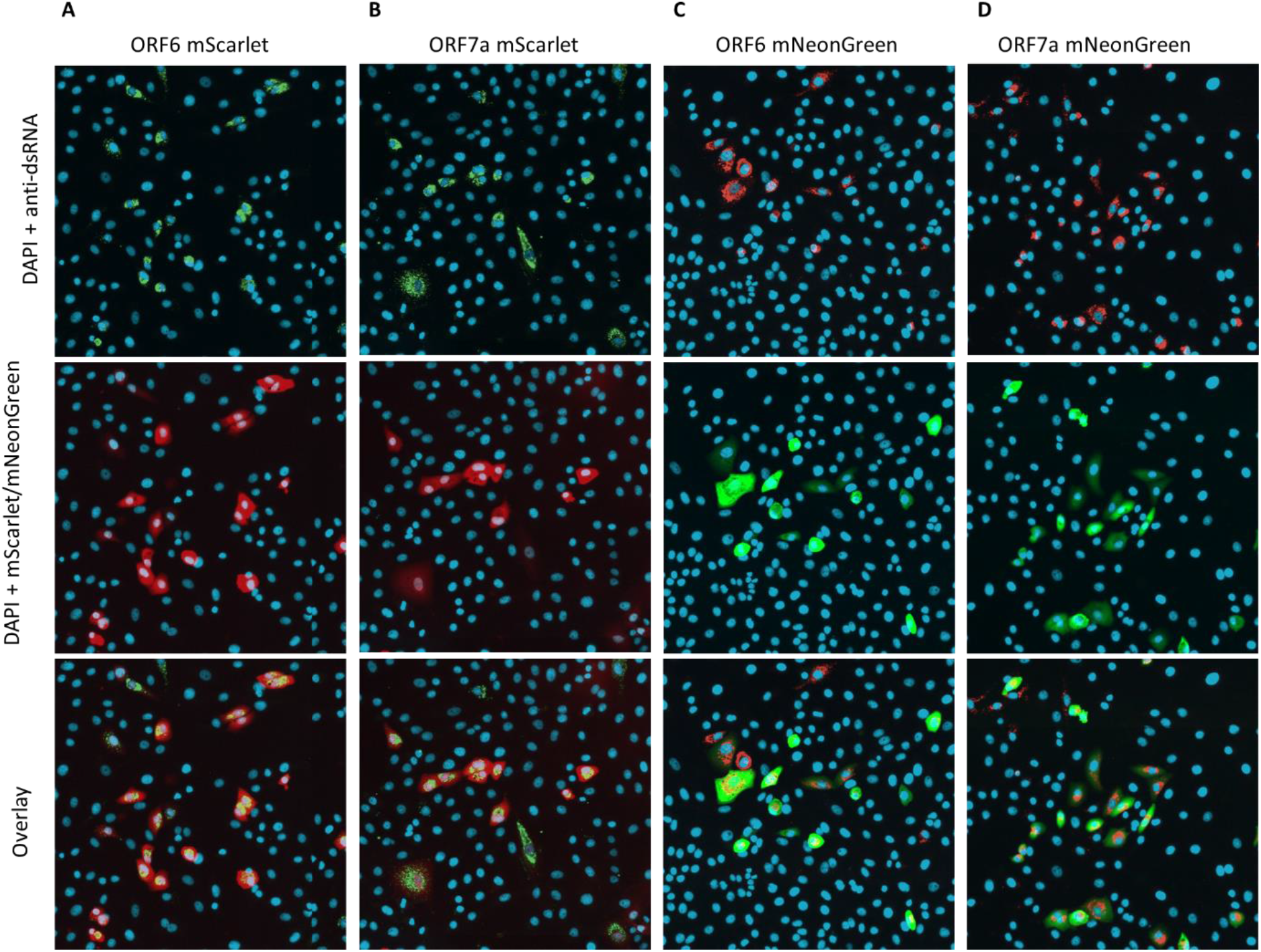
Detection of mScarlet, mNeonGreen and dsRNA after transfection of *in vitro* RNA transcripts derived from dual reporter replicons. VAT cells were transfected with *in vitro* RNA transcripts produced from A) pSC2-Rep-Wu-p-6Scar, B) pSC2-Rep-Wu-p-7Scar, C) pSC2-Rep-Wu-p-6NG and D) pSC2-Rep-Wu-p-7NG replicon constructs and an N gene construct. Cells in the top panels were stained for dsRNA and nuclear DNA (DAPI). The middle panels show fluorescence from the mScarlet and mNeonGreen proteins and DAPI staining of nuclear DNA whilst the bottom panels show a three-colour overlay. Fluorescence from the reporter proteins was acquired in the TRITC channel (mScarlet) or the FITC channel (mNeonGreen) using an ImageExpressPico microscope.

*In vitro* RNA transcripts produced from the four replicon constructs were then tested for their ability to mediate RLuc expression. VAT cells were transfected with *in vitro* RNA transcripts from the four replicon constructs and assayed for RLuc activity at 24-, 36- and 48-hours post-transfection (Figure 5). The kinetics of RLuc expression resulting from the different replicon *in vitro* RNA transcripts was similar to that observed previously using the pSC2-Rep-Wu-Gp-RL replicon *in vitro* RNA transcripts, with maximal expression at 36 hours post-transfection in the VAT cells. The level of peak expression was also of a similar magnitude as before (ranging from 1.8 – 4.5 × 10^7^ RLU). There were some differences in RLuc expression resulting from transfection of the different replicon *in vitro* RNA transcripts, which could have been due to differences in the quality of the transcripts, the transfection efficiency or expression levels from the ORF6 or ORF7 sgRNAs encoding the mNeonGreen/mScarlet fluorophores. As for the pSC2-Rep-Wu-Gp-RL replicon, numerous attempts were made to isolate cell populations stably harbouring all four of the dual reporter replicons by puromycin selection. Selection was attempted using a range of cell types, but as for the pSC2-Rep-Wu-Gp-RL replicon, stable cell line selection was not possible, suggesting that replicon replication was still cytotoxic despite the presence of the nsp1 mutations.

**Figure 5.**
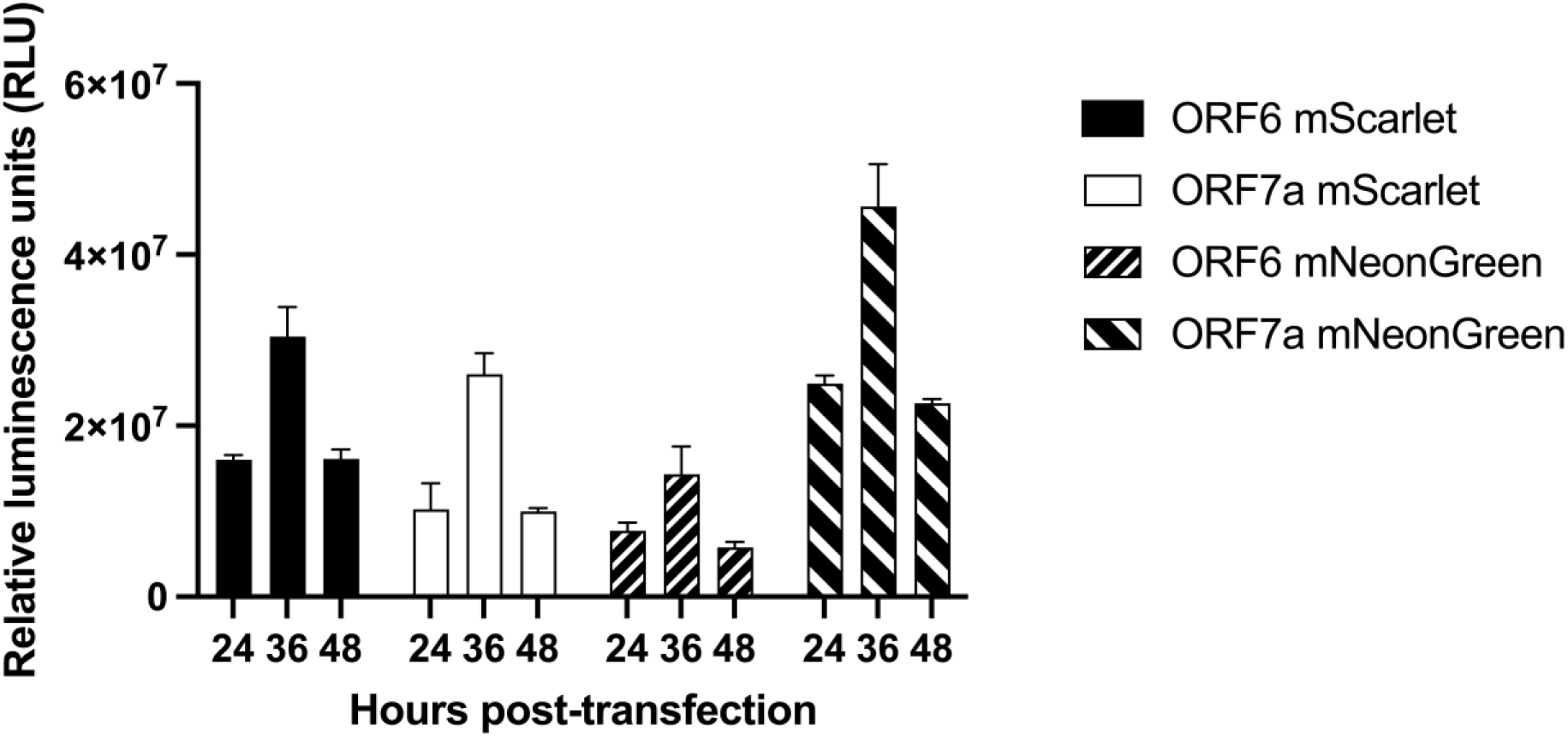
Rluc activity after transfection of *in vitro* RNA transcripts derived from dual reporter replicons. VAT cells were transfected with *in vitro* RNA transcripts from the dual reporter replicons pSC2-Rep-Wu-p-RL-6Scar (ORF6 mScarlet), pSC2-Rep-Wu-p-RL-7Scar (ORF7 mScarlet), pSC2-Rep-Wu-p-RL-6NG (ORF6 mNeonGreen), pSC2-Rep-Wu-p-RL-7NG (ORF7 mNeonGreen) and an N gene construct. The sample luminescence was adjusted for assay background by subtracting the cell only control. Graphs show mean and standard deviation of technical triplicates.

### Construction of a dual-reporter replicon for the SARS-CoV-2 VOC Delta

As the SARS-CoV-2 pandemic has progressed new VOCs have emerged, carrying numerous lineage defining mutations warranting study. In addition, the mutations present in different VOCs may influence virus cytotoxicity. At the time of writing this manuscript, only one other study has reported the development of a replicon system derived from a VOC genome. In this case only the S coding sequence was deleted to allow the assembly of VLPs (42). A replicon corresponding to the SARS-CoV-2 VOC Delta was therefore produced with the continued aim of isolating cells stably harbouring a SARS-CoV-2 replicon. The same design was adopted as for the dual reporter replicons pSC2-Rep-Wu-p-RL-6Scar and pSC2-Rep-Wu-p-RL-7NG, as they were shown to produce *in vitro* RNA transcripts which mediated expression of either mScarlet/mNeonGreen and gave the highest levels of RLuc expression in VAT cells. The S and M gene sequences were replaced with the *pac* and *Rluc* gene coding sequences respectively and the ORF6 and ORF7a coding sequences with those of mScarlet and mNeonGreen respectively (Figure 1B). Lineage defining mutations characteristic of the SARS-CoV-2 VOC Delta were introduced into the replicon during the assembly process by substitution of cDNA fragments corresponding to the Wuhan sequence with those produced by RT-PCR amplification from SARS-CoV-2 VOC Delta genomic RNA. In addition, OL-PCR and Hi-Fi assembly were used to engineer the mNeonGreen/mScarlet reporter genes into the VOC Delta genetic background. The fragments were then assembled into BAC/YAC plasmids to produce the replicon constructs pSC2-Rep-Del-p-RL-6Scar and pSC2-Rep-Del-p-RL-7NG.

Caco2/ACE2 and VAT cells were transfected with *in vitro* RNA transcripts derived from the pSC2-Rep-Del-p-RL-6Scar and pSC2-Rep-Del-p-RL-7NG replicons. A proportion of the cells were fixed at 24 hours post-transfection and examined for the presence of dsRNA, the N protein and the expression of the introduced fluorophore, with the remainder of the cells harvested at 24-, 48-, 72- and 96-hours post-transfection for the examination of RLuc activity (Figure 6). The results showed that *in vitro* transcripts derived from both replicons were capable of mediating replicon replication, as evidenced by the presence of dsRNA and the expression of the marker proteins from sgRNAs. The assay of Rluc activity in lysates from VAT transfected cells showed similar kinetics of Rluc expression as for the previous replicon constructs. As in previous experiments numerous attempts were made to isolate cell populations that stably maintained SARS-CoV-2 VOC Delta replicons under puromycin selection. However, after 10-15 days under selection no viable cells were obtained.

**Figure 6.**
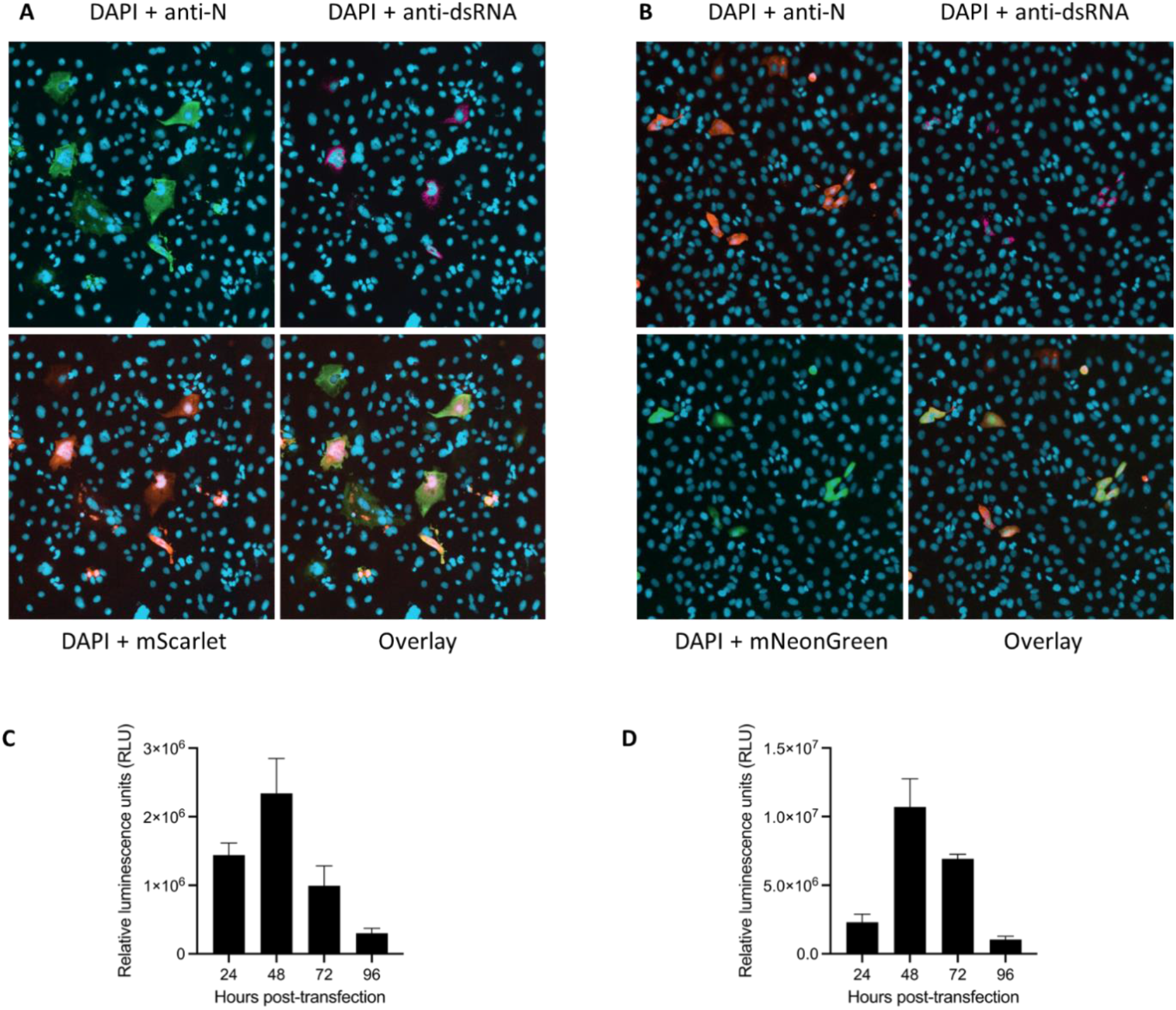
Validation of the replication of *in vitro* RNA transcripts derived from SARS-CoV-2 VOC Delta replicons. A and C) Caco2/ACE2 and B and D) VAT cells were transfected with *in vitro* RNA transcripts produced from pSC2-Rep-Del-p-RL-6Scar and pSC2-Rep-Del-p-RL-7NG respectively and an N gene construct. At 24 hours post-transfection a portion of the cells were fixed and stained with antibodies recognising the N protein and dsRNA and examined for fluorescence from A) mScarlet, B) mNeonGreen and DAPI stained nuclear DNA. The bottom right panels show a three-colour overlay. Fluorescence from the reporter proteins was acquired in the TRITC channel (mScarlet) or the FITC channel (mNeonGreen) using an ImageExpressPico microscope. Expression of RLuc in cells transfected with *in vitro* RNA transcripts derived from C) pSC2-Rep-Del-p-RL-6Scar and D) pSC2-Rep-Del-p-RL-7NG was monitored over the period 24-96 hours post-transfection. The sample luminescence was adjusted for assay background by subtracting the cell only control. Graphs show mean and standard deviation of n = 1 biological repeat performed as technical triplicates.

### Evaluation of the replicon system as an antiviral screening tool

Although it has not been possible to isolate cells stably expressing any of the replicon constructs produced to date, it was of interest to determine if the dual reporter replicons could faithfully be used for antiviral screening. Therefore, the replication of replicons derived from pSC2-Rep-Wu-p-RL-7NG and pSC-Rep-Del-p-RL-7NG were tested for their sensitivity to remdesivir using VAT cells. VAT cells were transfected with *in vitro* RNA transcripts produced from pSC2-Rep-Wu-p-RL-7NG and pSC2-Rep-Del-p-RL-7NG. The cells were seeded into 96-well plates and then incubated with a half-log fold 8-point dilution series of remdesivir starting with 20 µM as the highest dose. After 24 hours of incubation, prior to the peak of maximal RLuc activity in previous assays, the cells were harvested, and cell lysates assayed for RLuc activity (Figure 7).

**Figure 7.**
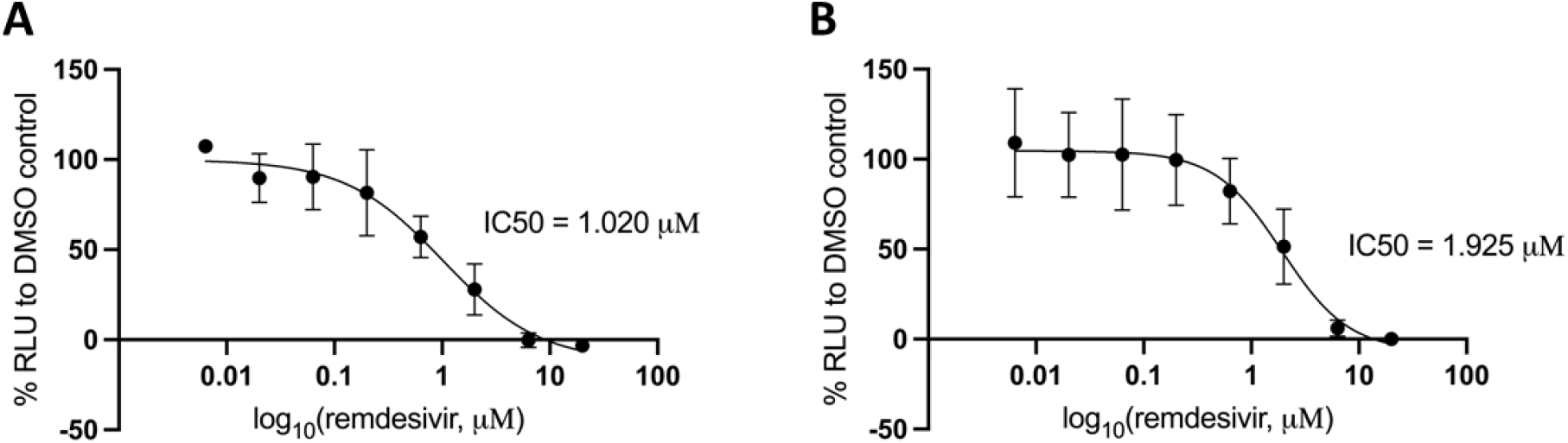
Effects of remdesivir on the replication of replicon RNA transcripts. VAT cells were transfected with *in vitro* RNA transcripts produced from an N gene construct and the A) pSC2-Rep-Del-p-RL-7NG and B) pSC2-Rep-Wu-p-RL-7NG replicons. The transfected cells were incubated with a half-log fold dilution series of remdesivir, starting at 20 µM. At 24 hours post-transfection the cells were lysed and assayed for luciferase activity. The graphs show mean and standard deviation as a percentage of the DMSO control, adjusted for background. Panel A shows n=3 independent biological repeats done in technical triplicate; panel B shows a technical triplicate. The assay background was determined from the RLuc signal produced after transfection of uncapped (mock) replicon *in vitro* RNA transcripts.

Using *in vitro* RNA transcripts from the Delta (pSC2-Rep-Del-p-RL-7NG) and Wuhan (pSC2-Rep-Wu-p-RL-7NG) replicons, IC50 values of 1.02 µM and 1.925 µM respectively were determined by fitting a nonlinear regression curve to the data. These values are in the range of IC50 values obtained for remdesivir using assays based on SARS-CoV-2 infection of Vero cells and other SARS-CoV-2 replicon studies using Vero cells (33-35, 37, 39, 59). The results demonstrate that the replicons provide a valuable platform for antiviral testing.

A major aim of this study was the production of a cell line stably maintaining a SARS-CoV-2 replicon. Although the replicons produced in this study are replication competent, as evidenced by the expression of reporter genes engineered in place of the ORFs encoded by the S, M, ORF6 and ORF7a sgRNAs, it was not possible to select cells stably maintaining the replicons, despite repeated attempts using different cell lines and conditions. A recent study describes the establishment of a SARS-CoV-2 replicon in BHK cells stably expressing the SARS-CoV-2 N protein. The nsp1 K164A/H165A mutation was introduced into the replicon and found necessary to produce stable replicon cell clones (37). Despite including these mutations it was still not possible to recover cell populations stably maintaining the replicons, as reported in another study (39). Spontaneous mutations in nsp4 and nsp10 were identified in almost all replicon RNA populations isolated for cells stabling maintaining the replicons, which could be suggestive of adaptive mutations that allowed for non-cytopathic replication in BHK-21 cells (37). Furthermore, a study reporting the isolation of Vero E6 populations stably maintaining a minimal replicon did not include the nsp1 mutations in the replicon genome, although the sequences of the stably maintained replicons were not determined (59).

Transfection of different cell lines revealed that VAT cells were highly permissive for replicon transfection, with transfection efficiencies between 10-20%. Caco2 cells were the most efficiently transfected human cell line with a transfection efficiency of ~5%. Huh 7.5, A549-A2T2 and VTN cells showed a transfection efficiency of ~1-3%. Both VAT and VTN cells were produced from VeroE6 cells; suggesting that cell specific differences in replicon transfection efficiency existed in the original cells used for producing the VAT and VTN cells or differences arose during the selection and propagation of the cells. It would be interesting to investigate whether the VAT cell line promotes increased SARS-CoV-2 replicon replication as they have been found to increase virus production (46).

The Wuhan-Hu-1 and Delta VOC dual-reporter replicons established in this investigation provide a useful and convenient tool for transient gene expression studies with read-out from two reporter proteins. Furthermore, the finding that remdesivir inhibits replicon replication demonstrates that the replicons can be used for drug screening and will be useful for dissecting the mechanism of action of drugs. As the replicons lack the S coding sequence they provide a useful tool to study the effects of drugs on intracellular virus replication. The results of this study also show that the initially developed replicon system can be adapted to introduce VOC lineage defining mutations with relative ease.

## Funding and Acknowledgments

The work was supported by UK Research and Innovation / Medical Research Council grants MR/V027506/1 and MR/R020566/1 to ADD and MR/W005611/1/ to ADD and DAM who are also supported by the United States Food and Drug Administration (contract 5F40120C00085). MKW and TJ were in part supported by grant MR/R020566/1. This work was supported by funding to ADD from the Elizabeth Blackwell Institute, funded in part by the Wellcome Trust (Grant number 794 204813/Z/16/Z). The authors would like to acknowledge support of the University of Bristol’s Alumni and Friends, which funded the ImageXpress Pico Imaging System.

## Author contributions

ME and ADD conceived/designed the study, ME, MKW, TJ, JB, DAM and ADD generated molecular reagents and protocols used in the study, ME generated and analysed the data; ME and ADD interpreted the data; ADD and DAM acquired funding for the study. ME and ADD prepared the manuscript. All authors provided critical review of the manuscript.

